# Study comparing characteristics of ademetionine-containing tablets from different countries

**DOI:** 10.64898/2026.03.27.714742

**Authors:** José M. Mato, Grace L. H. Wong, Yiscka Gooijer, Azadeh Safaei

## Abstract

**Background/Objectives:** The quality and characteristics of approved medicines can vary substantially depending on manufacturing processes and standards within a given country. The aim of the study was to compare the available marketed brands of ademetionine tablets derived from various countries in order to identify potential differences between the different formulations.

**Methods:** We performed comprehensive analyses of the physical, chemical, and dissolution characteristics of different formulations of ademetionine tablets marketed in China, India, Russia, Ukraine, and Uzbekistan, using the originator formulation of Heptral® as the reference standard. The formulations were evaluated at initial analysis and after 3 months at 40°C/75% relative humidity. Clinical parameters such as ademetionine content, degradation products, S,S-isomer, and water content were assessed using HPLC, and a dissolution profile analysis performed in 2 hours of acid solution followed by 90 minutes in a buffer solution.

**Results:** The Nusam (India) and Ximeixin (China) products were the two products most comparable to the Heptral products. Adenomak (Ukraine), the only food-grade product and only one with the tosylate salt showed the most significant quality variations compared to Heptral including dissolution failure as well as considerable variability between batches.

**Conclusions:** The study highlights the importance of using pharmaceutical-grade ademetionine products to maintain clinical efficacy and ensuring standards are maintained across global markets.

## Introduction

Ademetionine, also known as S-adenosylmethionine (SAMe), is a compound found in virtually all tissues and body fluids, playing a key role in numerous biochemical processes including methylation reactions, epigenetic regulation, detoxification reactions, phospholipid synthesis and glutathione syntheses [1-6]. It is synthesized from methionine and ATP by methionine adenosyltransferase (MAT) in the methionine cycle and serves as the main methyl donor in one-carbon metabolism, supporting thousands of reactions catalyzed by methyltransferase including DNA and histone methylation, phosphatidylcholine synthesis, and neurotransmitter regulation [1,3]. After donating its methyl group, ademetionine is converted to S-adenosylhomocysteine (SAH), linking it to homocysteine metabolism and the transsulfuration pathway that leads to synthesis of cysteine, taurine (involved in bile acid conjugation) and the major cellular antioxidant glutathione (required for hepatic detoxification) [1,2,4,6,7]. Through aminopropylation, ademetionine is also a precursor to polyamine synthesis, predominantly spermidine and spermine, which play a crucial role in regulation of transcription and translation, cell growth and apoptosis [5].

Ademetionine has two chiral centers a chiral carbon at the α-amino position, and a sulfonium sulfur [8]. The chiral sulfur center exists in two enantiomeric forms, (S,S)- and (R,S)-ademetionine [8]. The (S,S) configuration is the only biosynthesized form and is the biologically active form for almost all ademetionine dependent methyltransferases [8]. In its native form, ademetionine is labile and degrades rapidly, and thus for pharmaceutical development it is in a stable salt form [3].

Ademetionine has been used in the treatment of liver disorders, depression, osteoarthritis, fibromyalgia and fatigue [1,3,9-11]. The formulation of a medication can be as important as its efficacy in determining how successful a treatment is. It can play a critical role in absorption, distribution, and elimination of a drug, which in turn can influence the clinical profile of a medication and have a significant impact on patient quality of life, disease outcomes, and adherence to the treatment protocol [12,13]. For ademetionine, increased degradation or dissolution may reduce its clinical effects. The different salt forms used in different formulations may also affect the stability, bioavailability, and ultimately, the clinical efficacy of the product. Thus, some preparations on the market may differ or be superior to others with regard to stability and quality [14]. For example, one study found that a novel formulation (MSI-195) had a significantly higher bioavailability compared with a commercially available product, SAM-e Complete [15]. Several salts of ademetionine with strong organic or inorganic acids, including polyacids, are known, however, only sulfuric acid, 1,4-butanedisulfonic acid, and p-toluenesulfonic acid (tosylated) salts are presently on the market [7]. The active pharmaceutical ingredient of ademetionine formulation (Heptral®) manufactured by Abbott Products Operations AG is ademetionine 1,4-butanedisulfonate.

The aim of this study was to compare the available marketed brands of ademetionine tablets derived from various countries in order to identify potential differences between the different formulations in terms of physical attributes, chemical attributes, and drug release parameters. Analysis was performed on Heptral® 400 and 500 mg tablets, together with generic ademetionine tablets using multiple batches with varying shelf-life based on market availability. Many of the generic ademetionine tablets in the countries in this study have a similar composition of ademetionine ion, although some have recently changed from food products to drug products and/or have changed the ademetionine salt used in the formulation from tosylate to 1,4-butanedisulfonate.

## Methods

### Study design

Physical and chemical characteristics of 400 and 500 mg ademetionine tablets from different brands marketed in China, India, Russia, Ukraine, and Uzbekistan were evaluated. Multiple batches with varying shelf-life were used based on market availability. The products were studied at release (T=0; initial analysis) and after 3 months in a commercial pack (primary packaging material) at accelerated conditions of 40°C/75% relative humidity (T=3M) to assess stability. The in-house specification of Heptral tablets were considered as a reference for the comparison.

### Physical and chemical parameters evaluated

#### Physical parameters

At T=0, tablet dimensions (thickness, length, and width) were measured according to European Pharmacopoeia standards, tablets and packaging were examined visually, and the surface and internal topology of the tablets were examined. Uniformity of weight was assessed according to European Pharmacopoeia standards at both T=0 and T=3M, with average weight calculated based on the available number of tablets.

#### Chemical parameters

Chemical parameters were analyzed using high pressure liquid chromatography (HPLC), hydrophilic interaction liquid chromatography (HILIC) principle, with ultraviolet (UV) detection. Ademetionine content (mg/tablet and % relative label claim) was determined at T=0 and T=3M, and a stability slope (mg/tablet/month) was created. Degradation products ADN (adenine), SAH (S-adenosyl-L-homocysteine), MTA (methylthioadenosine), DecaSaMe (decarboxylated ademetionine), and SAMe-G (SAMe-glycin isomer-2) and impurities along with unspecified and total impurities (% relative label claim) were also assessed at T=0 and T=3M, and stability slopes (% relative label claim/month) were created. Degradation products homoserine (HS) and 2-amino-4-butyrolactone (ABL) (% relative label claim) were assessed at T=0 and T=3M, and stability slopes (% relative label claim/month) were created. As the HS and ABL impurities do not have a chromophore, these are not visible using the HPLC technique used for the other degradation products and thus HS and ABL were analyzed using a separate HPLC with UV detection method. The methods utilized for the assay, degradation product and HS/ABL analysis were in-house Abbott methods, which are fully validated for the Heptral products and, by definition, for the competitor drugs. The diasterioisomeric form of SAMe (S,S-isomer) and water content (min-max and average) were analyzed by HPLC with UV detection and Karl Fischer titration, respectively, at T=0 and T=3M, and stability slopes created (% min-max/month). For all parameters, stability slopes were compared to those for Heptral, with similar stability resulting in a ratio equal to 1.0.

Finally, a dissolution profile analysis was conducted to check release in 0.1N hydrochloric acid (HCL) for two hours (acid stage) followed by the buffer stage in pH 6.8 (quality control [QC] media) on six tablets per batch at T= 5, 10, 15,20, 30, 45, and 90 minutes. The dissolution analysis was performed using a dissolution robot with autosampler, sample filtration over 1 µm GXP filter, flow through cuvette (0.02 cm) and on-line (UV) detector, aligned with the European Pharmacopoeia 2.9.3 Dissolution Test for Solid Dosage Forms.

## Results

A summary of the characteristics of the analyzed batches of ademetionine is presented in **Table 1**. The shelf-life of the products varied between 14 months and 36 months. The batches were studied at different points in their shelf-life, with initial testing of all batches within the shelf-life apart from one batch of Samelix 400 (461021) which was 5 months past the expiry date at initial analysis. All batches apart from the Adenomak products from Ukraine were formulated using 1,4-butanedisulphonic acid. Adenomak was the only product that was food grade. In terms of packaging, most of the preparations were available and provided in aluminum packaging.

**Table 1:** Characteristics of the ademetionine gastro-resistance tablet brands analyzed in the study.

| Strength (mg) | Brand name | Manufacturer | Market | Batch numbers | Shelf-life | Shelf-life at initial analysis (April 2024) | Salt | Pack | Prescription required |
| --- | --- | --- | --- | --- | --- | --- | --- | --- | --- |
| 400 | Heptral | Abbott | Global | 1194211 | 36 months | 11 months | 1,4-butanedisulphonic acid | Alu – Alu blisters | No |
|  | Nusam | Sun Pharma laboratories Ltd. | India | CPA0071<br>CPA0012 | 18 months<br>14 months | 10 months<br>14 months | 1,4-butanedisulphonic acid | Alu – Alu blisters | Yes |
|  | Cirrosam | La Renon Healthcare Private limited | India | D223188<br>D223290 | 18 months<br>18 months | 13 months<br>11 months | 1,4-butanedisulphonic acid | Alu – Alu blisters | Yes |
|  | Samelix | Kanopharma | Russia | 461021<br>260822<br>030223 | 24 months<br>24 months<br>24 months | 29 months<br>20 months<br>14 months | 1,4-butanedisulphonic acid | PVC/PE/PVDC-Alu blisters | Yes |
|  | Ademetionine | Velpharm | Russia | 390722<br>600623 | 36 months<br>36 months | 21 months<br>13 months | 1,4-butanedisulphonic acid | PVC/PE/PVDC-Alu blisters | No |
|  | Ademta | Dr. Sertus | Uzbekistan | 11101513<br>208004360 | 36 months<br>36 months | 29 months<br>20 months | 1,4-butanedisulphonic acid | Alu – Alu blisters | Yes |
| 500 | Heptral | Abbott | Global | 1188470 | 36 months | 12 months | 1,4-butanedisulphonic acid | Alu – Alu blisters | Yes |
|  | Ximeixin | Zhejiang Hisun Pharmaceutical | China | 22303164 | 24 months | 13 months | 1,4-butanedisulphonic acid | Alu – Alu blisters | Not stated |
|  | Samelix | Kanopharma | Russia | 010323 | 24 months | 13 months | 1,4-butanedisulphonic acid | Alu – Alu blisters | Yes |
|  | Hepamethion | Arterium | Ukraine | 0075626<br>0063551<br>0069005 | 24 months<br>24 months<br>24 months | 12 months<br>20 months<br>17 months | 1,4-butanedisulphonic acid | Alu – Alu blisters | Yes |
|  | Adenomak | Macleods | Ukraine | KAY2301A<br>KAY2219A<br>KAY2225A | 24 months<br>24 months<br>24 months | 12 months<br>8 months<br>16 months | Tosylate | Alu – Alu blisters | Not a medicine product. Food product for special medical purposes |

Alu, aluminum; PE, polyethylene; PVDC, polyvinylidene chloride; PVC, polyvinyl chloride.

### Assessment of physical parameters

A summary of the physical characteristics of the batches of ademetionine to Heptral is presented in **Table 2**. In terms of surface coating, Nusam, Ademta and Adenomak are the only products comparable to Heptral. The ademetionine (Russia) and Hepamethion products have no gastro-resistant coating which may allow degradation of the product, including the active ingredient. There were notable differences in weight between the products with Cirrosam and Samelix 400mg being the heaviest compared to Heptral 400 mg and Adenomak notably heavier than Heptral 500 mg.

**Table 2:** Comparison of the physical characteristics of ademetionine products compared with Heptral.

|  | Batch numbers | Tablet thickness, mm | Nature of surface coating | Thickness of surface coating | Gastro-resistant coating confirmed<sup>a</sup> | Average weight, mg<sup>b</sup> |
| --- | --- | --- | --- | --- | --- | --- |
| 400 mg |  |  |  |  |  |  |
| Heptral | 1194211 | 6.9 | - | - | Yes | 920.7 |
| Nusam | CPA0071 | 6.6 | Comparable | Comparable | Yes | 931.9 |
|  | CPA0012 | 6.6 |  |  |  | 924.1 |
| Cirrosam | D223188 | 7.0 | Different, more compact | Comparable | Yes | 1124.9 |
|  | D223290 | 7.0 |  |  |  | 1130.7 |
| Samelix | 461021 | 6.0 | Different, more compact | Comparable | Yes | 1009.6 |
|  | 260822 | 5.8 |  |  |  | 1007.8 |
|  | 030223 | 5.9 |  |  |  | 992.9 |
| Ademetionine (Russia) | 390722 | 5.8 | Different, more compact | Comparable | No | 958.4 |
|  | 600623 | 5.6 |  |  |  | 952.6 |
| Ademta | 11101513 | 7.9 | Comparable | Comparable | Yes | 973.1 |
|  | 208004360 | 7.3 |  |  |  | 961.5 |
| 500 mg |  |  |  |  |  |  |
| Heptral | 1188470 | 6.9 | - | - | Yes | 1163.5 |
| Ximeixin | 22303164 | 6.4 | Different | Comparable | Yes | 1131.2 |
| Samelix | 010323 | 7.8 | Different, more compact | Slightly thinner | Yes | 1289.2 |
| Hepamethion | 0075626 | 6.7 | Different, more compact | Approx. twice as thin | No | 1125.9 |
|  | 0063551 | 6.7 |  |  |  | 1151.1 |
|  | 0069005 | 6.8 |  |  |  | 1154.8 |
| Adenomak | KAY2301A | 6.4 | Comparable | Comparable | Not stated | 1476.9 |
|  | KAY2219A | 6.3 |  |  |  | 1472.0 |
|  | KAY2225A | 6.4 |  |  |  | 1473.6 |

<sup>a</sup>The absence of a coating film can cause degradation of the tablet components, including the active ingredient, which may decrease the clinical efficacy of the product.
<sup>b</sup>The average weight was assessed based on a sample of 12 tablets for all batches.

### Assessment of chemical parameters

A summary of the chemical characteristics of the ademetionine batches compared with Heptral is presented in **Tables 3–6**.

**Table 3:**
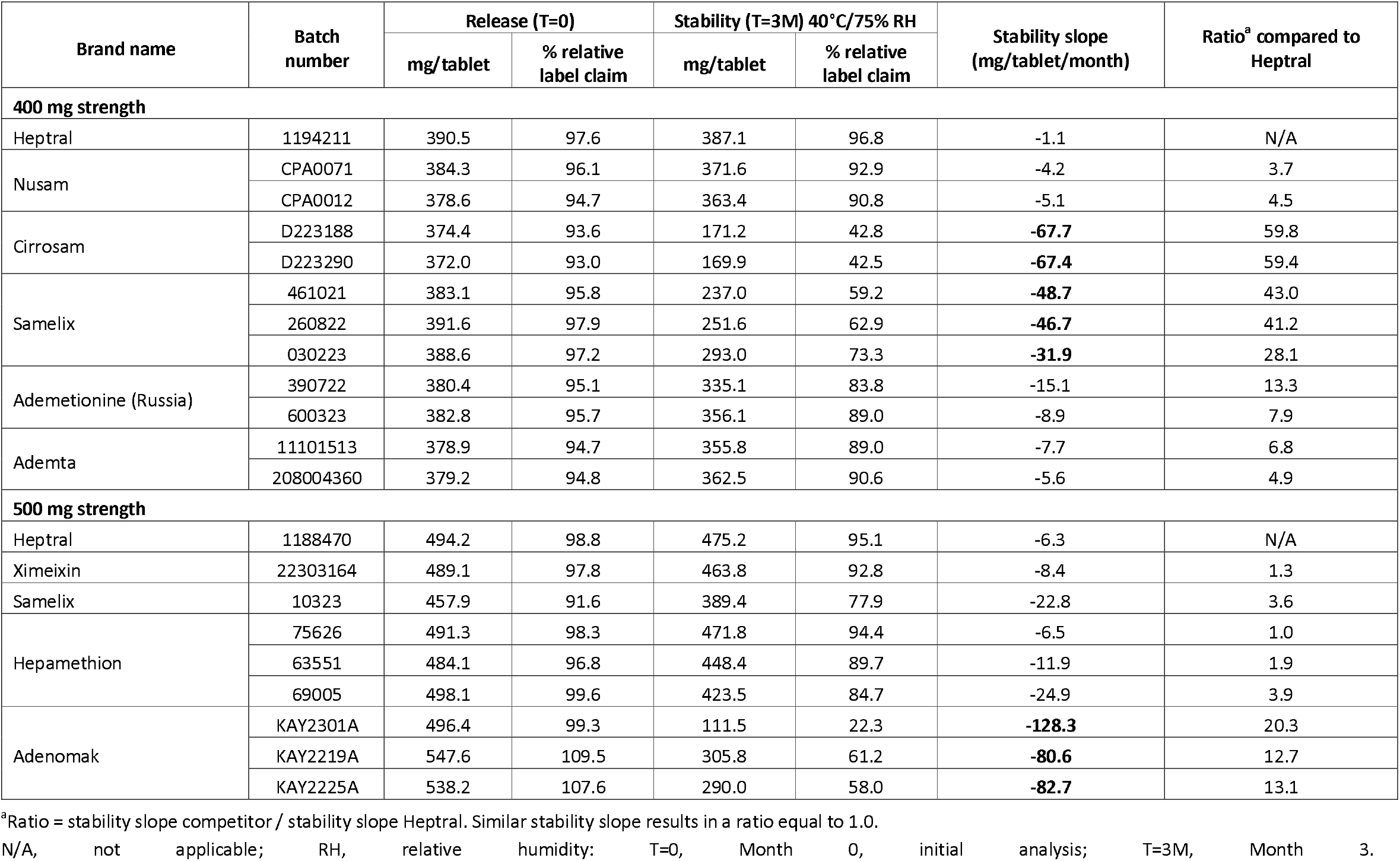
Ademetionine content.

**Table 4:** Degradation products/related substances release and stability (% relative label claim)

| Brand name | Batch number | Release (T=0)<br>(% relative label claim) |  |  |  |  |  |  |  | Stability (T=3M) 40°C / 75% RH<br>(% relative label claim) |  |  |  |  |  |  |  |
| --- | --- | --- | --- | --- | --- | --- | --- | --- | --- | --- | --- | --- | --- | --- | --- | --- | --- |
|  |  | MTA | ADN | Deca-SAMe | SAMe-G | SAH | Largest unknown (single) | Total unknown | Total degradation<sup>a</sup> | MTA | ADN | Deca-SAMe | SAMe-G | SAH | Largest Unknown (single) | Total unknown | Total degradation<sup>a</sup> |
| 400 mg strength |  |  |  |  |  |  |  |  |  |  |  |  |  |  |  |  |  |
| Heptral | 1194211 | 0.52 | 0.29 | 0.20 | 0.06 | 0.17 | BPQL | BPQL | 1.54 | 0.80 | 0.69 | 0.19 | 0.09 | 0.47 | 0.06 | 0.06 | 2.76 |
| Nusam | CPA0071 | 0.55 | 0.51 | N.D. | 0.07 | 0.32 | BPQL | BPQL | 1.69 | 0.71 | 0.95 | N.D. | 0.17 | 0.59 | 0.08 | 0.08 | 2.93 |
|  | CPA0012 | 0.78 | 0.33 | 0.98 | N.D. | 0.35 | 0.06 | 0.06 | 3.27 | 1.05 | 0.64 | 1.00 | N.D. | 0.66 | 0.11 | 0.18 | 4.53 |
| Cirrosam | D223188 | 0.86 | 0.71 | N.D. | BPQL | 0.38 | 0.08 | 0.08 | 2.44 | 1.12 | 1.80 | N.D. | BPQL | 1.68 | 0.77 | 0.77 | 8.27 |
|  | D223290 | 0.88 | 0.72 | N.D. | BPQL | 0.39 | 0.07 | 0.07 | 2.48 | 0.65 | 2.38 | N.D. | BPQL | 1.54 | 0.84 | 0.84 | 8.30 |
| Samelix | 461021 | 0.48 | 1.06 | N.D. | BPQL | 0.36 | 0.06 | 0.06 | 2.49 | 0.20 | 7.90 | N.D. | N.D. | 0.82 | 0.37 | 1.64 | 12.45 |
|  | 260822 | 0.39 | 0.94 | N.D. | N.D. | 0.24 | BPQL | BPQL | 1.84 | 0.20 | 7.81 | N.D. | N.D. | 0.71 | 0.27 | 1.40 | 11.46 |
|  | 030223 | 0.44 | 0.82 | N.D. | BPQL | 0.37 | BPQL | BPQL | 1.93 | 0.29 | 5.90 | N.D. | BPQL | 0.80 | 0.27 | 0.83 | 8.94 |
| Ademetionine (Russia) | 390722 | 0.41 | 0.77 | N.D. | BPQL | 0.28 | BPQL | BPQL | 2.13 | 0.36 | 3.44 | N.D. | BPQL | 0.67 | 0.15 | 0.28 | 5.68 |
|  | 600323 | 0.29 | 0.47 | N.D. | BPQL | 0.15 | BPQL | BPQL | 1.95 | 0.37 | 1.99 | N.D. | 0.13 | 0.44 | 0.06 | 0.06 | 3.66 |
| Ademta | 11101513 | 0.48 | 1.18 | N.D. | BPQL | 0.43 | 0.06 | 0.06 | 2.60 | 0.55 | 2.20 | N.D. | N.D. | 0.72 | 0.12 | 0.17 | 4.28 |
|  | 208004360 | 0.71 | 0.74 | N.D. | BPQL | 0.40 | 0.07 | 0.06 | 2.34 | 0.78 | 1.74 | N.D. | 0.10 | 0.80 | 0.14 | 0.14 | 4.55 |
| Specifications (%) |  | ≤2.5 | ≤2.5% | N/A | N/A | ≤2.0 | ≤0.20 | ≤3.0 | ≤6.0 | ≤2.5 | ≤2.5 | N/A | N/A | ≤2.0 | ≤0.20 | ≤3.0 | ≤6.0 |
| 500 mg strength |  |  |  |  |  |  |  |  |  |  |  |  |  |  |  |  |  |
| Heptral | 1188470 | 0.57 | 0.26 | 0.18 | 0.13 | 0.17 | BPQL | BPQL | 1.54 | 0.82 | 0.64 | 0.18 | 0.13 | 0.47 | 0.06 | 0.06 | 2.79 |

CONFIDENTIAL 27 March 2026
| Brand name | Batch number | Release (T=0)<br>(% relative label claim) |  |  |  |  |  |  |  | Stability (T=3M) 40°C / 75% RH<br>(% relative label claim) |  |  |  |  |  |  |  |
| --- | --- | --- | --- | --- | --- | --- | --- | --- | --- | --- | --- | --- | --- | --- | --- | --- | --- |
|  |  | MTA | ADN | Deca-SaMe | SaMe-G | SAH | Largest unknown (single) | Total unknown | Total degradation<sup>a</sup> | MTA | ADN | Deca-SaMe | SaMe-G | SAH | Largest Unknown (single) | Total unknown | Total degradation<sup>a</sup> |
| Ximeixin | 22303164 | 0.35 | 0.29 | 0.13 | BPQL | 0.18 | BPQL | BPQL | 1.33 | 0.69 | 0.86 | 0.11 | 0.00 | 0.5 | 0.06 | 0.06 | 2.77 |
| Samelix | 010323 | 0.45 | 2.07 | N.D. | BPQL | 0.44 | 0.06 | 0.06 | 3.31 | 0.27 | 4.94 | N.D. | BPQL | 0.66 | 0.17 | 0.33 | 6.97 |
| Hepamethion | 75626 | 0.31 | 0.34 | 0.24 | N.D. | 0.28 | BPQL | BPQL | 1.41 | 0.45 | 1.29 | 0.26 | 0.08 | 0.54 | 0.07 | 0.11 | 3.09 |
|  | 63551 | 0.35 | 0.39 | 0.25 | N.D. | 0.30 | BPQL | BPQL | 1.41 | 0.53 | 1.42 | 0.25 | 0.08 | 0.60 | 0.07 | 0.14 | 3.39 |
|  | 69005 | 0.29 | 0.38 | 0.27 | BPQL | 0.23 | BPQL | BPQL | 1.40 | 0.41 | 1.19 | 0.26 | 0.08 | 0.45 | 0.06 | 0.06 | 2.85 |
| Adenomak | KAY2301A | 0.81 | 2.02 | N.D. | BPQL | 0.53 | 0.15 | 0.27 | 4.04 | 0.30 | 9.10 | N.D. | BPQL | 0.96 | 0.89 | 4.39 | 21.17 |
|  | KAY2219A | 0.72 | 0.82 | N.D. | BPQL | 0.42 | 0.06 | 0.11 | 2.29 | 0.69 | 8.31 | N.D. | BPQL | 1.80 | 0.66 | 2.31 | 17.46 |
|  | KAY2225A | 0.93 | 0.72 | N.D. | BPQL | 0.38 | 0.06 | 0.12 | 2.40 | 1.01 | 7.91 | N.D. | BPQL | 2.19 | 0.83 | 2.26 | 18.65 |
| Specifications (%) |  | ≤2.5 | ≤2.5 | ≤0.8 | ≤0.5 | ≤2.0 | ≤0.20 | ≤3.0 | ≤6.0 | ≤2.5 | ≤2.5 | ≤0.8 | ≤0.5 | ≤2.0 | ≤0.20 | ≤3.0 | ≤6.0 |

<sup>a</sup>Total degradation is the sum of the related substances reported (HS/ABL).
ADN, adenine; BPQL, below practical quantification limit; DecaSaMe, decarboxylated ademethionine; MTA, methylthioadenosine; N/A, not applicable; ND, not determined; RH, relative humidity; SAH, S-adenosyl-L-homocysteine; SaMe-G, SaMe-glycine isomer-2; T=0, Month 0, initial analysis; T=3M, Month 3.

**Table 5.** Homoserine and 2-amino-4-butyrolactone release and stability (% relative label claim)

| Brand name | Batch number | Release (T=0)<br>(% relative label claim) |  | Stability (T=3M) 40°C / 75% RH<br>(% relative label claim) |  | Stability slope<br>(% relative label claim / month) |  | Ratio<sup>a</sup> compared to Heptral |  |
| --- | --- | --- | --- | --- | --- | --- | --- | --- | --- |
|  |  | HS | ABL | HS | ABL | HS | ABL | HS | ABL |
| 400 mg strength |  |  |  |  |  |  |  |  |  |
| Heptral | 1194211 | BPQL | 0.06 | 0.31 | 0.39 | N/A | 0.03 | N/A | N/A |
| Nusam | CPA0071 | BPQL | 0.05 | 0.25 | 0.38 | N/A | 0.04 | N/A | 1.6 |
|  | CPA0012 | 0.15 | 0.18 | 0.62 | 0.82 | 0.01 | 0.07 | N/A | 2.5 |
| Cirrosam | D223188 | 0.10 | 0.82 | 0.31 | 2.08 | 0.24 | 0.59 | N/A | 22.1 |
|  | D223290 | 0.10 | 0.82 | 0.32 | 2.07 | 0.24 | 0.58 | N/A | 21.9 |
| Samelix | 461021 | 0.19 | 0.79 | 0.34 | 1.09 | 0.20 | 0.25 | N/A | 9.4 |
|  | 260822 | 0.10 | 0.52 | 0.17 | 0.81 | 0.14 | 0.21 | N/A | 8.0 |
|  | 030223 | 0.09 | 0.42 | 0.21 | 0.66 | 0.11 | 0.15 | N/A | 5.6 |
| Ademetionine<br>(Russia) | 390722 | 0.19 | 0.33 | 0.47 | 0.60 | 0.05 | 0.04 | N/A | 1.6 |
|  | 600323 | 0.41 | 0.20 | 0.63 | 0.47 | -0.07 | -0.05 | N/A | -2.0 |
| Ademta | 11101513 | 0.13 | 0.20 | 0.32 | 0.43 | 0.02 | 0.04 | N/A | 1.4 |
|  | 208004360 | 0.15 | 0.35 | 0.28 | 0.63 | 0.07 | 0.12 | N/A | 4.4 |
| Specifications (%) |  | ≤ 1.2 | ≤ 1.2 | ≤ 1.2 | ≤ 1.2 | N/A |  |  |  |
| 500 mg strength |  |  |  |  |  |  |  |  |  |
| Heptral | 1188470 | BPQL | 0.06 | 0.23 | 0.43 | N/A | 0.07 | N/A | N/A |
| Ximeixin | 22303164 | 0.09 | 0.09 | 0.29 | 0.47 | 0.00 | 0.06 | N/A | 0.9 |
| Samelix | 10323 | 0.08 | 0.25 | 0.2 | 0.52 | 0.06 | 0.11 | N/A | 1.6 |
| Hepamethion | 75626 | 0.06 | 0.07 | 0.18 | 0.28 | 0.00 | 0.03 | N/A | 0.5 |
|  | 63551 | BPQL | 0.08 | 0.11 | 0.29 | N/A | 0.06 | N/A | 0.9 |
|  | 69005 | 0.07 | 0.09 | 0.16 | 0.32 | 0.01 | 0.05 | N/A | 0.8 |
| Adenomak | KAY2301A | N.D. | 2.29 | 0.41 | 4.14 | N/A | 1.24 | N/A | 18.7 |
|  | KAY2219A | N.D. | 2.16 | 0.21 | 2.19 | N/A | 0.66 | N/A | 9.9 |

27 March 2026
| Brand name | Batch number | Release (T=0)<br>(% relative label claim) |  | Stability (T=3M) 40°C / 75% RH<br>(% relative label claim) |  | Stability slope<br>(% relative label claim / month) |  | Ratio<sup>a</sup> compared to Heptral |  |
| --- | --- | --- | --- | --- | --- | --- | --- | --- | --- |
|  |  | HS | ABL | HS | ABL | HS | ABL | HS | ABL |
|  | KAY2225A | N.D. | 2.57 | 0.25 | 2.72 | N/A | 0.82 | N/A | 12.4 |
| Specifications (%) |  | ≤ 1.2 | ≤ 1.2 | ≤ 1.2 | ≤ 1.2 | N/A |  |  |  |

<sup>a</sup>Ratio = stability slope competitor / stability slope Heptral. Similar stability slope results in a ratio equal to 1.0.
ABL, 2-amino-4-butyrolactone; BPQL, below practical quantification limit; HS, homoserine N/A, not applicable; ND, not determined; RH, relative humidity; T=0, Month 0, initial analysis; T=3M, Month 3.

**Table 6.**
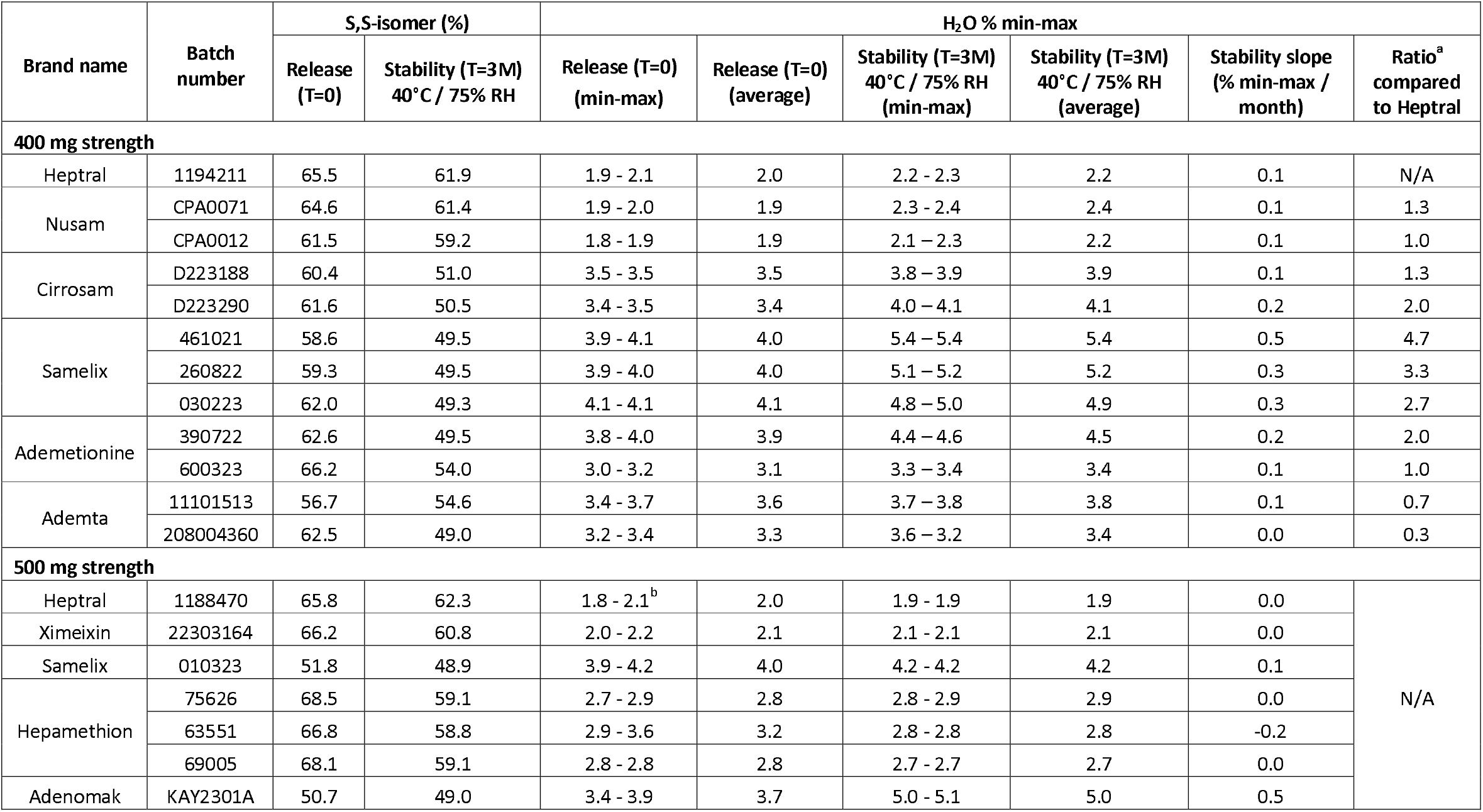
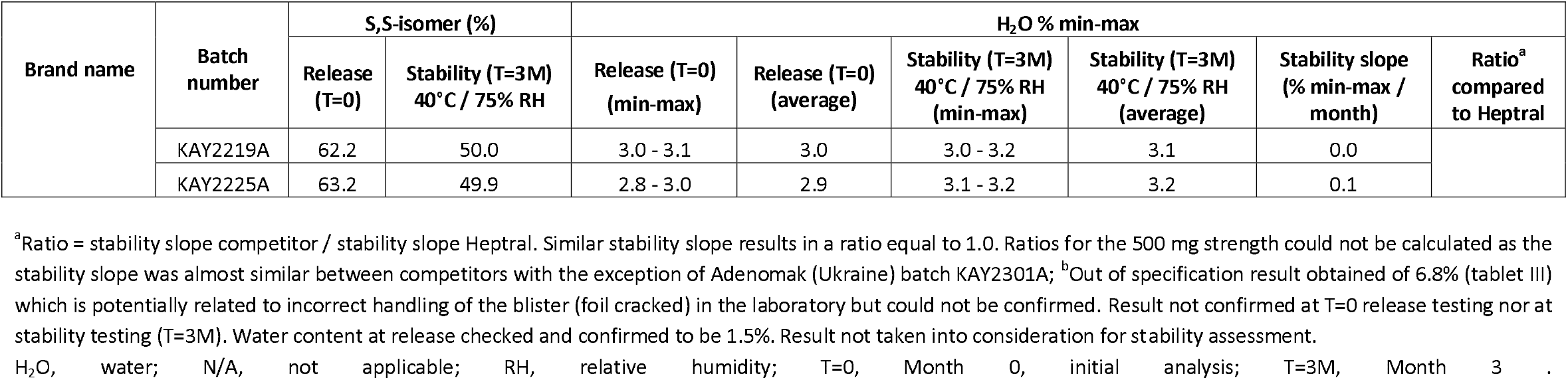
Ademetionine S,S-isomer and water content.

The Nusam (400mg, from India) and Ximeixin (500mg, from China) products were comparable in specifications to the Heptral products (400 mg and 500 mg, respectively), both at T=0 and T=3M (**Tables 3-5**).

In terms of ademetionine content, all products met the specifications for ademetionine content (assay) except for Adenomak (two of the three batches tested higher for ademetionine content [KAY2219A and KAY2225A] (**Table 3**). All batches of Adenomak, Samelix and Cirrosam did not meet the specification for total degradation (**Table 4**). The Adenomak products showed considerable variability between batches in the results for degradation products, and the largest greatest total degradation (**Tables 4 and 5**).

All products met specifications for S,S-isomer in the initial testing (T=0) except for Samelix 500mg, which was out of range (lower). The Adenomak products showed considerable variability in results at T=0 with one batch out of range (KAY2301A), but at T=3M, all batches were lower for S,S-isomer (**Table 6**).

In terms of stability slopes, Cirrosam and Samelix 400 mg showed the largest deviations compared to the Heptral with very marked reductions in stability slope, as did Adenomak compared to Heptral 500 mg (**Table 3**). A decline in the stability slope was observed for all products except for one batch of Hepamethion (75626), which was comparable to Heptral. There was variability between the three batches of Hepamethion in terms of stability slope.

The water content in the Nusam and Ximeixin brands were comparable to the Heptral products, while all other products showed a higher water content which was not compliant with the shelf-life specification of Heptral (**Table 6**).

Apart from Nusam and Ximeixin, which were comparable in specifications to the Heptral products, all other products showed faster degradation compared with Heptral at T=3M. In the dissolution rate assays, Nusam, Samelix (400 mg and 500 mg), Ximeixin, and Hepamethion underwent full dissolution within 45 minutes; and Cirrosam, ademetionine (Russia), and Ademta within 90 minutes. Samelix 400 mg showed high variance between the three batches with both faster and slower dissolution observed. For all three batches of Adenomak, the tablets became adhesive and stuck to the dissolution vessel after being exposed to the dissolution media, with no disintegration visible after 3 hours (including the 90 minutes in buffer stage) (**Figure 1; Supplementary Table 1**).

**Figure 1.**
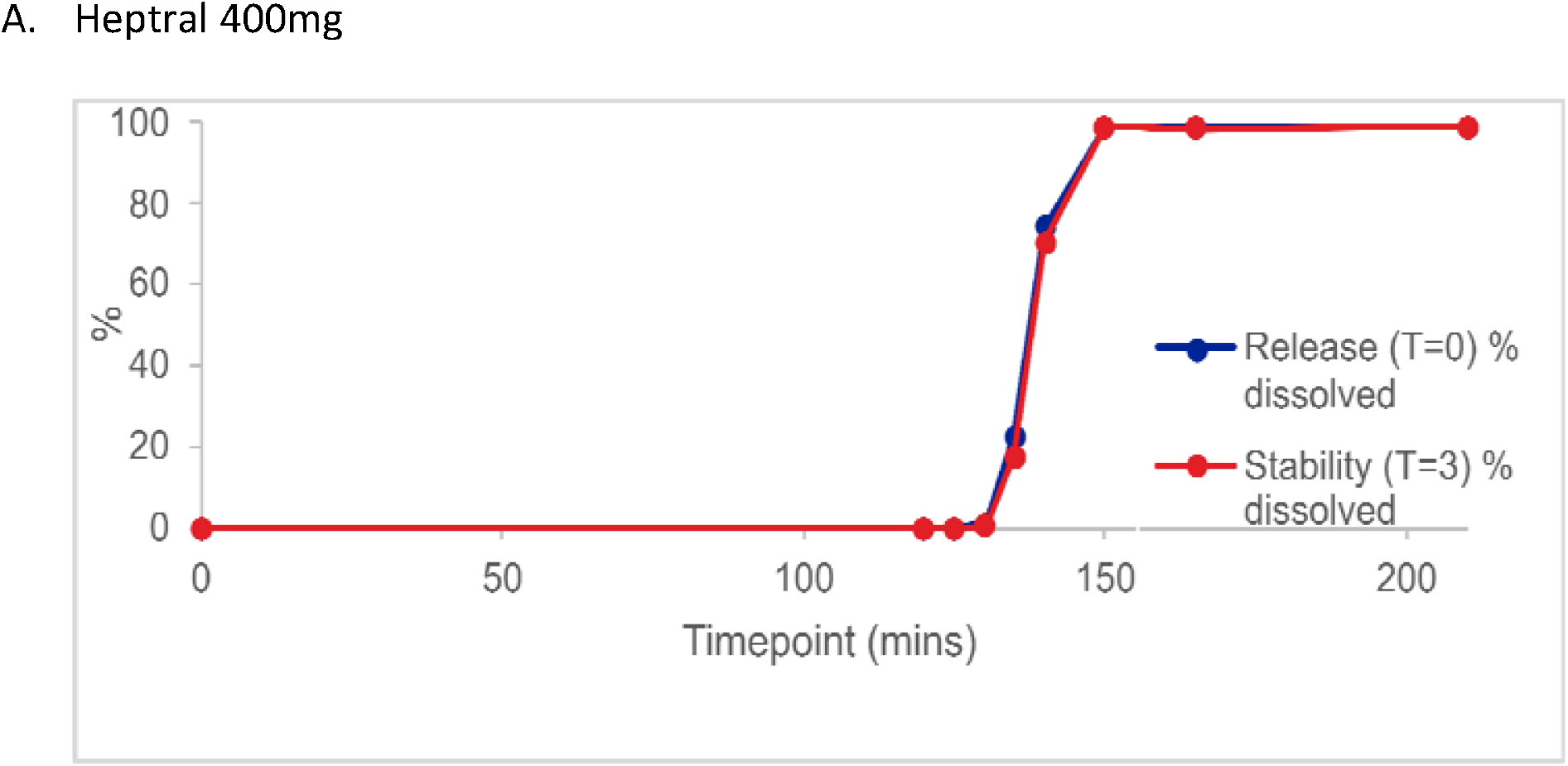
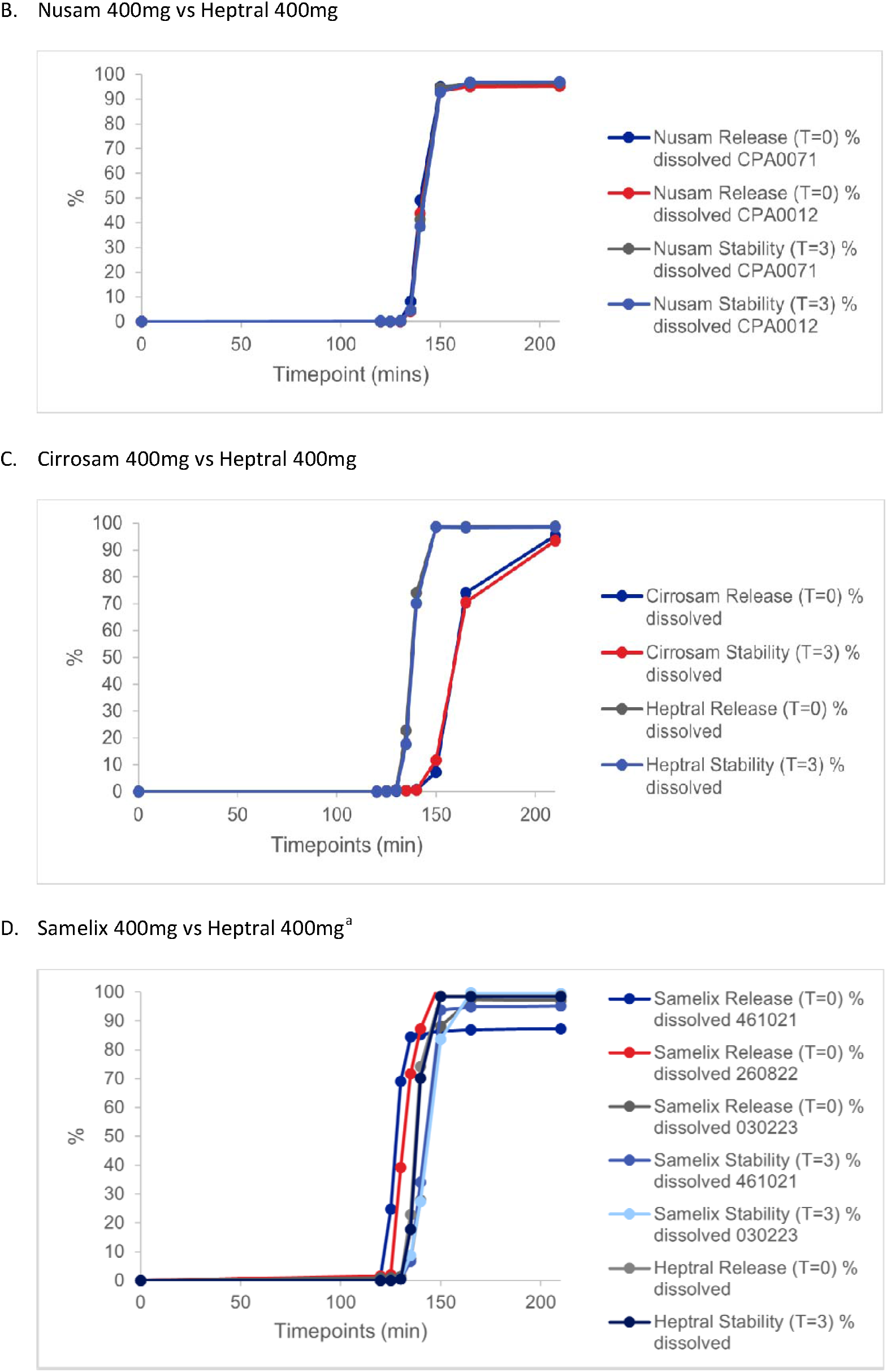
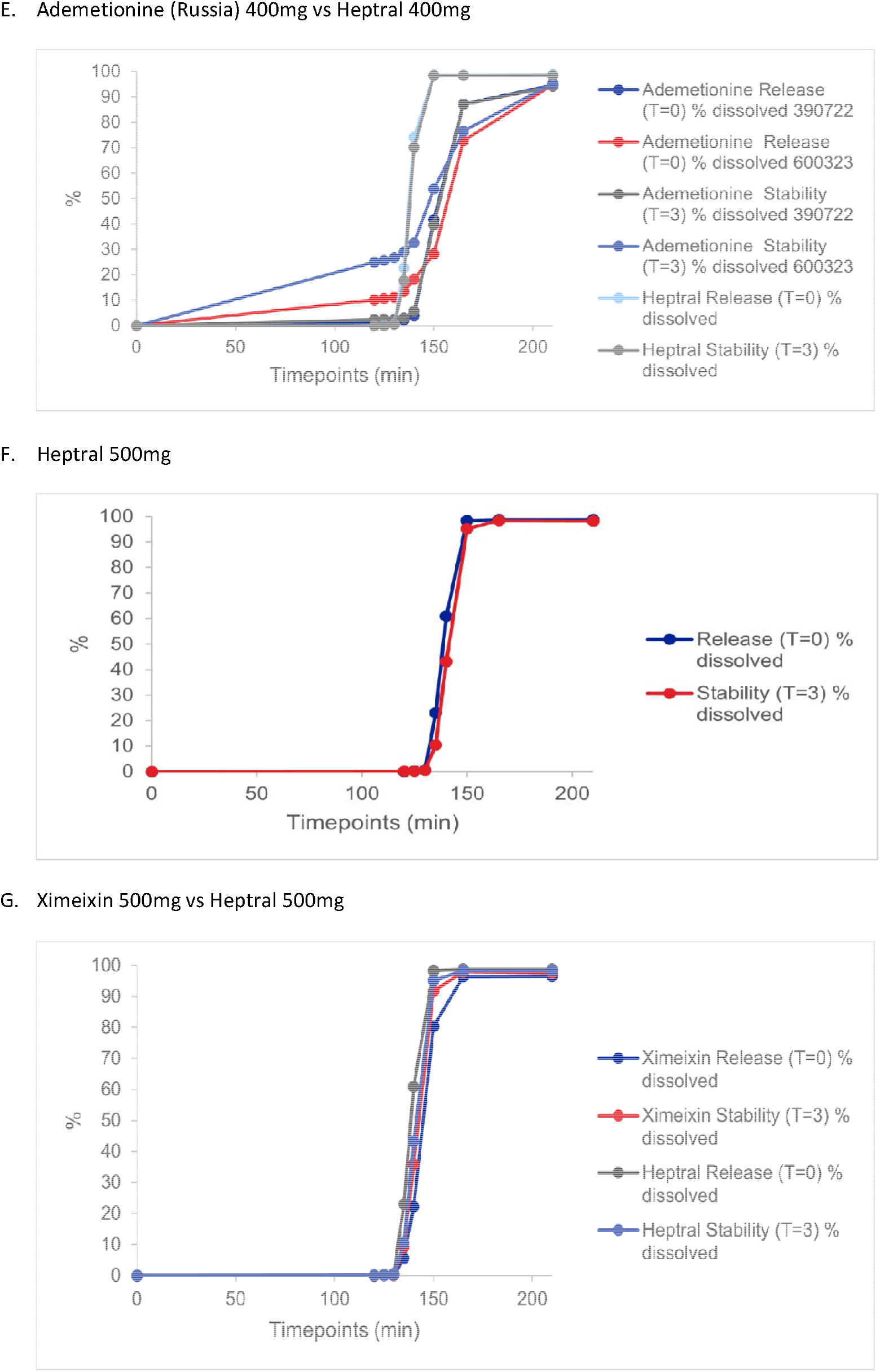
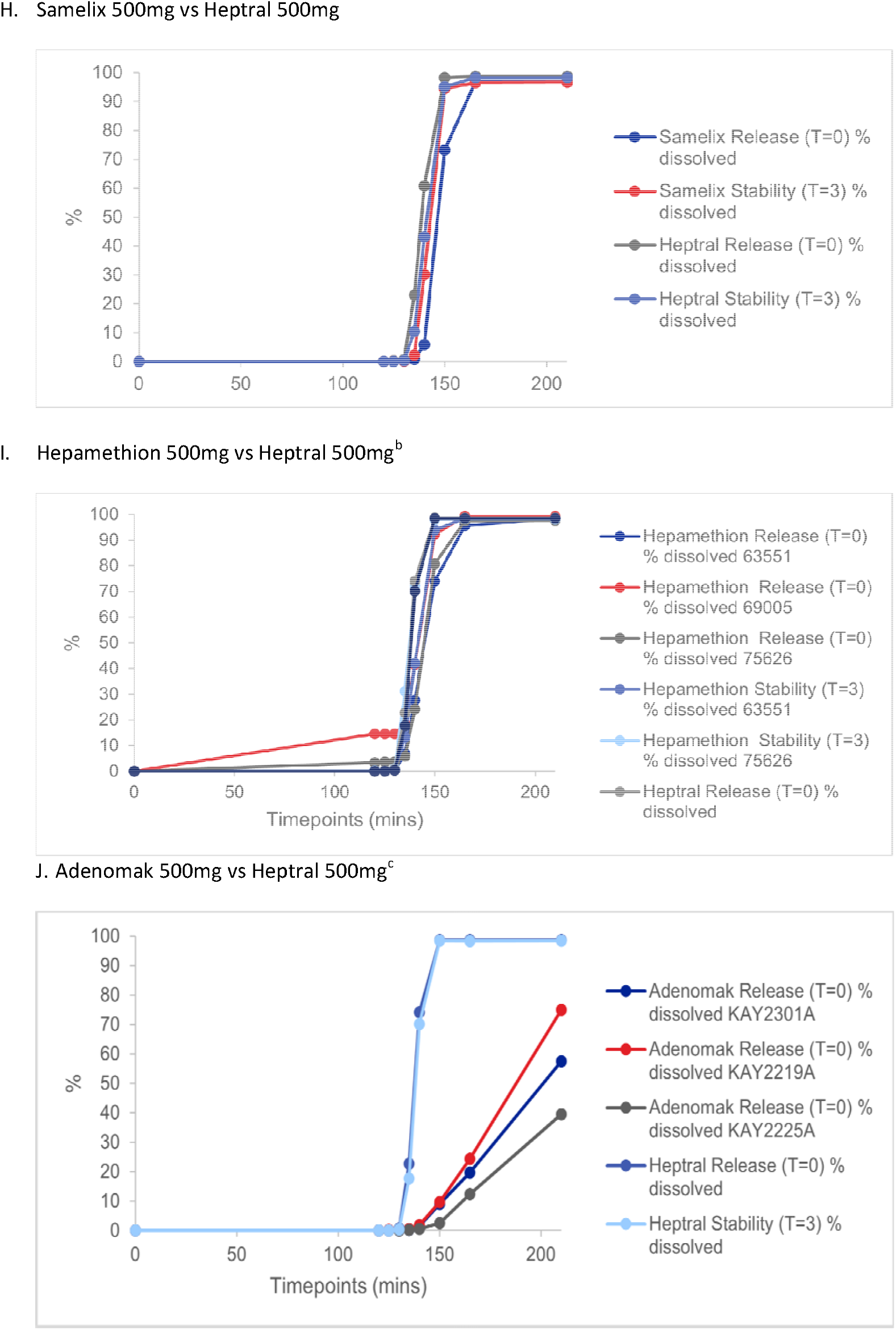
Dissolution profiles: Release at T=0 and stability at T=3 40°C/75% RH compared to Heptral.

The graphs show the timepoint from the beginning of the dissolution time: timepoints 0-120mins are two hours of acid stage where ≤ 10% is expected to have dissolved after 2 hours. Timepoints between 120 to 210 are the buffer stages which are tested at T= 0, 5, 10, 15,20, 30, 45, and 90 minutes where ≥ 75% within 90 min is expected. ^a^Stability data for Samelix 400mg Batch number 260822 was not assessed due to limited sample availability. ^b^Stability data for Hepamethion 500mg batch number 69005 is not available owing to interruption of the dissolution run due to instrument failure, and limited availability meaning no additional tablets available to repeat the analysis; ^c^No stability data for any of the Adenomak 500 mg batches as the tablets became adhesive and stuck to the dissolution vessel after being exposed to the dissolution media, with no disintegration visible after 210 min/90 min in buffer stage.

## Discussion

The study identified notable inter-country variation among 400 mg and 500 mg ademetionine tablet brands from China, India, Russia, Ukraine, and Uzbekistan relative to Heptral. Nusam (400mg product from India) and Ximeixin (500mg product from China) were the most similar in terms of physical and chemical properties to Heptral. While all pharmaceutical drug products undergo testing, the standards and regulations vary between different countries [16]. This may account for the differences in physical and chemical properties observed between the drug products tested from the different countries, however, the clinical significance of these differences is unknown.

The most notable variations in quality were observed for the only food-grade product tested, Adenomak from Ukraine. As well as more of the tested parameters (ademetionine content, degradation products and S,S isomer at 3 months) having specifications outside those of the Heptral products, there was also considerable variability between batches. All other products met the specifications for ademetionine content, degradation products and S,S-isomer in the initial testing except for Samelix 500mg which had lower S,S-isomer levels at initial testing.

Nusan and Ximeixin were the only products that had comparable water content to the Heptral products, all other products showed a higher water content which was not compliant with the shelf-life specification of Heptral.

The notable variations in quality with Adenomak are most likely due to the different regulatory standards applied between food and pharmaceutical drug products. Furthermore, Adenomak was the only product that contained the tosylate salt. In most cases, where a drug is marketed in more than one salt form, they are marketed as therapeutically equivalent and treated as identical products by clinicians. However, they may not be chemically equivalent [17]. The dissolution assay for Adenomak showed the most notable differences compared with the other products tested, with no dissolution seen over three hours. This could be due to the different salt formulation used in Adenomak compared with all the other products tested, suggesting that butanedisulfonate may be the superior salt formulation.

A reduction in the slope observed in stability studies—i.e., a slower decline in API concentration over time, can have far-reaching consequences on therapeutic efficacy and clinical outcomes. A decrease in this slope indicates more rapid API breakdown and thus a shorter effective shelf life. This accelerated degradation not only diminishes the available dose of active substance but also increases the risk of forming degradation products that may be pharmacologically inactive or potentially harmful. Clinically, such instability can lead to subtherapeutic dosing, orphaned dosing regimens, and unpredictable pharmacokinetic profiles, especially concerning when margin-of-error between therapeutic and toxic doses is narrow. From a formulation standpoint, any observed decrease in stability slope necessitates urgent reevaluation of excipient compatibility, container– closure system performance, and storage conditions. Ensuring optimal chemical stability through strategies preserve pharmacological activity, and ultimately safeguard patient efficacy and safety [18,19].

The packaging material did not appear to have an impact on the results. The ALU-ALU packing has been shown to offer superior protection against environmental factors like moisture, light, and oxygen compared with ALU-PVC, although it is costlier and less flexible in design [20].

The inter⍰brand differences observed in salt form (tosylate vs 1,4⍰butanedisulfonate), S,S⍰isomer content, water uptake, and delayed⍰release dissolution profiles are clinically relevant because they determine oral ademetionine exposure, with salt selection recognized as a critical determinant of solubility, stability, and therapeutic effect [21,22]. Under the Biopharmaceutics Classification System, inadequate enteric protection or slow/failed dissolution leads to bio⍰inequivalence even when label assay appears compliant, which can reduce treatment efficacy in liver disease or depression [23]. SAMe (ademetionine) is chemically labile and prone to degradation and epimerization of the active S,S⍰diastereomer in aqueous environments; therefore, formulation strategies that preserve the active isomer and ensure gastric bypass are essential for maintaining clinical effect [5,8].

Salt form selection is not a trivial matter: different salts can dramatically alter a drug’s solubility, dissolution rate, and subsequent bioavailability-critical pharmacokinetic parameters known to influence clinical outcomes. For other drugs, such as sodium valproate versus valproic acid, and glucosamine sulfate salts, clinical bioavailability differences have been documented between salt forms. Similarly, case studies in pharmaceuticals (e.g. ciprofloxacin and indinavir) demonstrate that salt-induced changes in dissolution rates directly affect the fraction absorbed (Fa) and therapeutic exposure [17,21,24-27].

In our study, Adenomak’s tosylate salt failed to dissolve over three hours, which suggests such a formulation could significantly underperform compared to the clinically validated butanedisulfonate form. This mirrors broader evidence showing that improper salt choice impairs absorption and decreases therapeutic effect [21,25].

More generally, variability in any of the tested formulation attributes—such as inadequate salt stabilization, low S,SIZisomer content, heightened water uptake, or subpar dissolution—can unpredictably impact systemic exposure, leading to diminished efficacy or increased variability in patient response. This is especially important for SAMe, which exhibits formulationIZdependent pharmacokinetics and a significant food effect, underscoring the need for tightly controlled product quality and administration conditions to optimize clinical outcomes [21,22].

Taken together, these findings support the preferential use of pharmaceutical⍰grade, enteric⍰coated ademetionine 1,4⍰butanedisulfonate tablets that meet compendial dissolution criteria, and caution against substituting salts or using dietary⍰supplement grade products without demonstrated equivalence, as variable exposure can compromise treatment effectiveness.

This study had a number of limitations to be considered when interpreting the results. First, the limited availability of tablets in some countries impacted the ability to carry out full analysis on some products. In addition, the shelf-life of the products at initial analysis varied between 8 and 29 months, which may have impacted the physical and chemical characteristics of the products. Finally, biological activity was not tested and therefore bioequivalence could not be assessed. As such, the likely clinical impact of the differences in physical and chemical properties is not known.

## Conclusions

Significant differences in quality attributes were found with the Adenomak brand (from Ukraine) compared with Heptral. Adenomak is the only food-grade product in the study, and the only one using the tosylate salt. These findings highlight the different regulatory standards between food and pharmaceutical drug products and emphasize the importance of ensuring the quality standards for medicines are adhered to and maintained across global markets.

## Supporting information

Heptral report Supplementary file

## Author contributions

Authors contributed equally to the conception, design, and writing of the manuscript. All authors critically revised the manuscript, agree to be fully accountable for ensuring the integrity and accuracy of the work, and read and approved the final manuscript.

## Data availability statement

Details regarding where data supporting reported results can be found, including links to publicly archived datasets analyzed or generated during the study. Where no new data were created, or where data is unavailable due to privacy or ethical restrictions, a statement is still required.

## Acknowledgments

Editorial assistance was provided by Martin Guppy at Metamols Limited.

## Abbreviations

The following abbreviations are used in this manuscript:

ABL: 2-amino-4-butyrolactone
AND: Adenine
ATP: Adenosine triphosphate DecaSaMe Decarboxylated ademethionine DNA Deoxyribonucleic acid
G: Glycine
HCL: Hydrochloric acid
HILIC: Hydrophilic interaction liquid chromatography
HPLC: High pressure liquid chromatography
HS: Homoserine
MAT: Methionine adenosyltransferase
MTA: Methylthioadenosine
QC: Quality control
RH: Relative humidity
SAH: S-adenosylhomocysteine
SAMe: S-adenosylmethionine
SAMe-G: SAMe-glycine isomer-2
UV: Ultraviolet

