## Supplementary material for "Study comparing characteristics of ademetionine-containing tablets from different countries": Heptral report Supplementary file

**Supplementary table 1:** **Dissolution profiles**

|  | **Timepoint (min)** | **Timepoint (min)  Buffer stage (pH 6.8)** | **Shelf-life specifications** | **Release (T=0)** | | | **Stability (T=3M) 40°C / 75% RH** | | | **Stability slope**  **(% dissolved/ month)** | **Ratio^b^**  **compared to Heptral** |
| --- | --- | --- | --- | --- | --- | --- | --- | --- | --- | --- | --- |
|  |  |  |  | **% dissolved** | **SD** | **% CV** | **% dissolved** | **SD** | **% CV** |  |  |
| Heptral 400mg (batch number: 1194211) | 0 | N/A | Acid Stage ≤ 10% dissolved after 2 hours  Buffer Stage Q ≥ 75% within 90 min^a^ | 0.0 | 0 | N/A | 0.0 | 0 | N/A | N/A | N/A |
|  | 120 | 0 |  | 0.0 | 0.0 | 155.7 | 0.0 | 0.0 | 172.5 |  |  |
|  | 125 | 5 |  | 0.1 | 0.0 | 40.4 | 0.0 | 0.0 | 106.1 |  |  |
|  | 130 | 10 |  | 0.7 | 0.9 | 134.6 | 0.4 | 0.4 | 89.7 |  |  |
|  | 135 | 15 |  | 22.8 | 8.3 | 36.2 | 17.7 | 7.7 | 43.3 |  |  |
|  | 140 | 20 |  | 74.1 | 9.6 | 13.0 | 70.1 | 9.2 | 13.1 |  |  |
|  | 150 | 30 |  | 98.6 | 2.2 | 2.3 | 98.5 | 1.6 | 1.6 |  |  |
|  | 165 | 45 |  | 98.6 | 2.2 | 2.3 | 98.4 | 1.6 | 1.6 | -0.1 |  |
|  | 210 | 90 |  | 98.7 | 2.2 | 2.2 | 98.5 | 1.6 | 1.6 | -0.1 |  |
| Nusam 400mg (batch number: CPA0071) | 0 | N/A | Acid Stage ≤ 10% dissolved after 2 hours  Buffer Stage Q ≥ 75% within 90 min^a^ | 0.0 | 0 | N/A | 0.0 | 0 | N/A | N/A | N/A |
|  | 120 | 0 |  | 0.0 | 0.0 | N/A | 0.0 | 0.0 | 224.3 |  |  |
|  | 125 | 5 |  | 0.1 | 0.0 | 33.3 | 0.1 | 0.1 | 51.6 |  |  |
|  | 130 | 10 |  | 0.1 | 0.1 | 74.1 | 0.2 | 0.2 | 87.8 |  |  |
|  | 135 | 15 |  | 8.2 | 7.0 | 85.2 | 4.6 | 3.0 | 66.0 |  |  |
|  | 140 | 20 |  | 49.0 | 16.2 | 33.1 | 41.3 | 7.9 | 19.0 |  |  |
|  | 150 | 30 |  | 94.8 | 2.7 | 2.9 | 94.5 | 2.0 | 2.1 |  |  |
|  | 165 | 45 |  | 96.1 | 2.5 | 2.6 | 96.3 | 2.3 | 2.4 | 0.1 | -1.0 |
|  | 210 | 90 |  | 96.3 | 2.6 | 2.7 | 96.6 | 2.4 | 2.5 | 0.1 | -1.5 |
| Nusam 400mg (batch number: CPA0012) | 0 | N/A | Acid Stage ≤ 10% dissolved after 2 hours  Buffer Stage Q ≥ 75% within 90 min^a^ | 0.0 | 0 | N/A | 0.0 | 0 | N/A | N/A | N/A |
|  | 120 | 0 |  | 0.0 | 0.0 | 187.7 | 0.2 | 0.2 | 90.7 |  |  |
|  | 125 | 5 |  | 0.1 | 0.1 | 128.9 | 0.0 | 0.0 | 158.4 |  |  |
|  | 130 | 10 |  | 0.0 | 0.0 | 175.4 | 0.3 | 0.3 | 120.5 |  |  |
|  | 135 | 15 |  | 4.2 | 3.1 | 75.5 | 4.6 | 5.4 | 118.8 |  |  |
|  | 140 | 20 |  | 43.8 | 12.4 | 28.3 | 38.5 | 11.3 | 29.4 |  |  |
|  | 150 | 30 |  | 93.1 | 3.8 | 4.1 | 92.7 | 3.7 | 4.0 |  |  |
|  | 165 | 45 |  | 95.1 | 2.1 | 2.2 | 96.8 | 2.0 | 2.1 | 0.6 | -8.5 |
|  | 210 | 90 |  | 95.2 | 2.1 | 2.3 | 96.9 | 2.1 | 2.2 | 0.6 | -8.5 |
| Cirrosam 400mg (batch number: D223188) | 0 | N/A | Acid Stage ≤ 10% dissolved after 2 hours  Buffer Stage Q ≥ 75% within 90 min^a^ | 0.0 | 0 | N/A | 0.0 | 0 | N/A | N/A | N/A |
|  | 120 | 0 |  | 0.0 | 0.0 | 244.9 | 0.0 | 0.1 | 244.9 |  |  |
|  | 125 | 5 |  | 0.1 | 0.0 | 48.6 | 0.1 | 0.1 | 87.2 |  |  |
|  | 130 | 10 |  | 0.1 | 0.1 | 62.5 | 0.2 | 0.1 | 70.6 |  |  |
|  | 135 | 15 |  | 0.3 | 0.1 | 45.4 | 0.0 | 0.1 | 243.5 |  |  |
|  | 140 | 20 |  | 0.6 | 0.5 | 86.6 | 0.2 | 0.2 | 99.3 |  |  |
|  | 150 | 30 |  | 10.8 | 12.8 | 118.2 | 7.5 | 9.3 | 124.4 |  |  |
|  | 165 | 45 |  | 71.8 | 11.9 | 16.6 | 67.7 | 13.3 | 19.6 | -1.4 | 20.5 |
|  | 210 | 90 |  | 96.7 | 3.1 | 3.2 | 96.1 | 2.7 | 2.8 | -0.2 | 3.0 |
| Cirrosam 400mg  **Batch Number:** D223290 | 0 | N/A | Acid Stage ≤ 10% dissolved after 2 hours  Buffer Stage Q ≥ 75% within 90 min^a^ | 0.0 | 0 | N/A | 0.0 | 0 | N/A | N/A | N/A |
|  | 120 | 0 |  | 0.0 | 0.0 | 79.3 | 0.0 | 0.0 | 169.8 |  |  |
|  | 125 | 5 |  | 0.1 | 0.1 | 41.1 | 0.1 | 0.1 | 83.8 |  |  |
|  | 130 | 10 |  | 0.2 | 0.1 | 27.3 | 0.3 | 0.1 | 36.8 |  |  |
|  | 135 | 15 |  | 0.4 | 0.1 | 27.7 | 0.4 | 0.2 | 48.2 |  |  |
|  | 140 | 20 |  | 0.7 | 0.5 | 62.6 | 0.6 | 0.2 | 38.7 |  |  |
|  | 150 | 30 |  | 7.2 | 5.1 | 71.0 | 11.7 | 8.0 | 68.0 |  |  |
|  | 165 | 45 |  | 74.0 | 4.1 | 5.5 | 70.5 | 15.0 | 21.3 | -1.2 | 17.5 |
|  | 210 | 90 |  | 95.4 | 3.5 | 3.7 | 93.4 | 2.2 | 2.4 | -0.7 | 10.0 |
| Samelix 400mg (batch number: 461021) | 0 | N/A | Acid Stage ≤10% dissolved after 2 hours  Buffer Stage Q ≥75% within 90 min^a^ | 0.0 | 0 | N/A | 0.0 | 0 | N/A | N/A | N/A |
|  | 120 | 0 |  | 1.2 | 0.6 | 49.6 | 0.0 | 0.0 | 116.3 |  |  |
|  | 125 | 5 |  | 24.8 | 14.5 | 58.5 | 0.2 | 0.2 | 82.3 |  |  |
|  | 130 | 10 |  | 69.0 | 4.6 | 6.6 | 0.5 | 0.1 | 26.3 |  |  |
|  | 135 | 15 |  | 84.4 | 3.8 | 4.5 | 6.7 | 1.9 | 28.5 |  |  |
|  | 140 | 20 |  | 85.2 | 3.8 | 4.4 | 34.1 | 12.2 | 35.7 |  |  |
|  | 150 | 30 |  | 86.5 | 3.9 | 4.5 | 93.8 | 1.9 | 2.0 |  |  |
|  | 165 | 45 |  | 87.0 | 3.7 | 4.2 | 94.9 | 1.4 | 1.5 | 2.6 | -39.5 |
|  | 210 | 90 |  | 87.3 | 3.6 | 4.2 | 95.1 | 1.3 | 1.4 | 2.6 | -39.0 |
| Samelix 400mg (batch number: 260822) | 0 | N/A | Acid Stage ≤10% dissolved after 2 hours  Buffer Stage Q ≥75% within 90 min^a^ | 0.0 | 0 | #DIV/0! | Not tested due to limited sample availability | | | N/A | N/A |
|  | 120 | 0 |  | 1.5 | 2.1 | 140.3 |  |  |  |  |  |
|  | 125 | 5 |  | 2.1 | 2.8 | 128.4 |  |  |  |  |  |
|  | 130 | 10 |  | 39.2 | 21.0 | 53.5 |  |  |  |  |  |
|  | 135 | 15 |  | 71.7 | 35.1 | 48.9 |  |  |  |  |  |
|  | 140 | 20 |  | 87.3 | 34.6 | 39.7 |  |  |  |  |  |
|  | 150 | 30 |  | 104.1 | 5.4 | 5.2 |  |  |  |  |  |
|  | 165 | 45 |  | 107.1 | 4.8 | 4.5 |  |  |  |  |  |
|  | 210 | 90 |  | 109.3 | 5.7 | 5.2 |  |  |  |  |  |
| Samelix 400mg (batch number: 030223) | 0 | N/A | Acid Stage ≤10% dissolved after 2 hours  Buffer Stage Q ≥75% within 90 min^a^ | 0.0 | 0 | N/A | 0.0 | 0 | N/A | N/A | N/A |
|  | 120 | 0 |  | 0.9 | 1.6 | 191.3 | 0.0 | 0.0 | 138.7 |  |  |
|  | 125 | 5 |  | 1.0 | 1.7 | 174.7 | 0.2 | 0.1 | 34.8 |  |  |
|  | 130 | 10 |  | 1.4 | 1.8 | 125.1 | 0.7 | 0.7 | 102.9 |  |  |
|  | 135 | 15 |  | 7.7 | 7.1 | 93.0 | 8.7 | 5.1 | 58.8 |  |  |
|  | 140 | 20 |  | 27.9 | 12.1 | 43.1 | 27.4 | 15.2 | 55.6 |  |  |
|  | 150 | 30 |  | 88.1 | 6.5 | 7.4 | 83.6 | 17.3 | 20.7 |  |  |
|  | 165 | 45 |  | 97.4 | 2.7 | 2.7 | 99.9 | 5.4 | 5.4 | 0.8 | -12.5 |
|  | 210 | 90 |  | 97.0 | 2.6 | 2.7 | 99.5 | 5.5 | 5.5 | 0.8 | -12.5 |
| Ademetionine 400mg (batch number: 390722) | 0 | N/A | Acid Stage ≤10% dissolved after 2 hours  Buffer Stage Q ≥75% within 90 min^a^ | 0.0 | 0 | N/A | 0.0 | 0.0 | N/A | N/A | N/A |
|  | 120 | 0 |  | 1.3 | 2.8 | 212.5 | 2.4 | 5.3 | 222.4 |  |  |
|  | 125 | 5 |  | 1.6 | 2.9 | 184.0 | 2.5 | 5.5 | 219.4 |  |  |
|  | 130 | 10 |  | 1.6 | 2.9 | 181.3 | 2.6 | 5.7 | 216.1 |  |  |
|  | 135 | 15 |  | 2.2 | 2.8 | 126.2 | 3.0 | 6.4 | 210.9 |  |  |
|  | 140 | 20 |  | 4.0 | 2.8 | 69.3 | 5.8 | 6.2 | 106.9 |  |  |
|  | 150 | 30 |  | 41.7 | 28.9 | 69.3 | 39.6 | 26.1 | 65.8 |  |  |
|  | 165 | 45 |  | 87.3 | 16.8 | 19.3 | 87.0 | 6.5 | 7.5 | -0.1 | 1.5 |
|  | 210 | 90 |  | 94.7 | 2.2 | 2.3 | 94.2 | 1.2 | 1.3 | -0.2 | 2.5 |
| Ademetionine 400mg (batch number: 600323) | 0 | N/A | Acid Stage ≤10% dissolved after 2 hours  Buffer Stage Q ≥75% within 90 min^a^ | 0.0 | 0 | N/A | 0.0 | 0 | N/A | N/A | N/A |
|  | 120 | 0 |  | 10.2 | 7.3 | 72.1 | 24.9 | 29.2 | 117.4 |  |  |
|  | 125 | 5 |  | 10.7 | 7.9 | 74.0 | 25.7 | 29.1 | 113.2 |  |  |
|  | 130 | 10 |  | 11.2 | 8.5 | 76.0 | 26.7 | 30.0 | 112.6 |  |  |
|  | 135 | 15 |  | 13.4 | 12.2 | 91.0 | 28.9 | 31.8 | 110.1 |  |  |
|  | 140 | 20 |  | 18.2 | 18.3 | 100.4 | 32.5 | 33.9 | 104.2 |  |  |
|  | 150 | 30 |  | 28.2 | 26.8 | 94.7 | 53.7 | 41.0 | 76.4 |  |  |
|  | 165 | 45 |  | 72.7 | 14.7 | 20.2 | 76.6 | 32.8 | 42.8 | 1.3 | -19.5 |
|  | 210 | 90 |  | 94.8 | 2.9 | 3.1 | 95.3 | 1.6 | 1.7 | 0.2 | -2.5 |
| Ademta 400mg (batch number: 11101513) | 0 | N/A | Acid Stage ≤10% dissolved after 2 hours  Buffer Stage Q ≥75% within 90 min^a^ | 0.0 | 0 | N/A | 0.0 | 0 | N/A | N/A | N/A |
|  | 120 | 0 |  | 0.0 | 0.0 | 0 | 0.0 | 0.0 | 78.6 |  |  |
|  | 125 | 5 |  | 0.0 | 0.0 | 137.5 | 0.1 | 0.0 | 32.4 |  |  |
|  | 130 | 10 |  | 0.1 | 0.0 | 38.1 | 0.1 | 0.1 | 45.5 |  |  |
|  | 135 | 15 |  | 3.2 | 3.7 | 115.9 | 3.0 | 4.1 | 136.8 |  |  |
|  | 140 | 20 |  | 25.3 | 22.6 | 89.3 | 22.4 | 16.9 | 75.1 |  |  |
|  | 150 | 30 |  | 87.3 | 10.0 | 11.5 | 86.4 | 14.8 | 17.1 |  |  |
|  | 165 | 45 |  | 94.9 | 2.1 | 2.2 | 98.1 | 2.6 | 2.7 | 1.1 | -16.0 |
|  | 210 | 90 |  | 94.9 | 2.2 | 2.3 | 98.2 | 3.0 | 3.1 | 1.1 | -16.5 |
| Ademta 400mg (batch number: 208004360) | 0 | N/A | Acid Stage ≤10% dissolved after 2 hours  Buffer Stage Q ≥75% within 90 min^a^ | 0.0 | 0 | N/A | 0.0 | 0 | N/A | N/A | N/A |
|  | 120 | 0 |  | 0.1 | 0.1 | 94.6 | 0.0 | 0.1 | 202.5 |  |  |
|  | 125 | 5 |  | 0.1 | 0.0 | 48.7 | 0.1 | 0.1 | 151.1 |  |  |
|  | 130 | 10 |  | 0.3 | 0.4 | 132.1 | 0.0 | 0.0 | 157.4 |  |  |
|  | 135 | 15 |  | 2.1 | 3.2 | 151.8 | 0.1 | 0.2 | 140.5 |  |  |
|  | 140 | 20 |  | 15.9 | 11.3 | 70.9 | 8.8 | 6.9 | 78.4 |  |  |
|  | 150 | 30 |  | 75.7 | 37.2 | 49.1 | 67.7 | 8.0 | 11.9 |  |  |
|  | 165 | 45 |  | 84.4 | 24.9 | 29.5 | 94.7 | 2.0 | 2.1 | 3.4 | -51.5 |
|  | 210 | 90 |  | 92.9 | 4.4 | 4.8 | 96.6 | 2.9 | 3.0 | 1.2 | -18.5 |
| **500 mg strength** | | | | | | | | | | | |
| Heptral  500 mg (batch number: 1188470) | 0 | N/A | Acid Stage ≤10% dissolved after 2 hours  Buffer Stage Q ≥75% within 90 min^a^ | 0.0 | 0 | N/A | 0.0 | 0 | N/A | N/A |  |
|  | 120 | 0 |  | 0.0 | 0.0 | 88.5 | 0.1 | 0.1 | 110.1 |  |  |
|  | 125 | 5 |  | 0.2 | 0.0 | 29.3 | 0.1 | 0.1 | 127.3 |  |  |
|  | 130 | 10 |  | 0.6 | 0.4 | 69.3 | 0.4 | 0.7 | 171.6 |  |  |
|  | 135 | 15 |  | 23.0 | 10.5 | 45.5 | 10.5 | 5.5 | 52.8 |  |  |
|  | 140 | 20 |  | 60.8 | 9.4 | 15.4 | 43.2 | 4.5 | 10.5 |  |  |
|  | 150 | 30 |  | 98.3 | 0.9 | 0.9 | 95.1 | 2.6 | 2.7 |  |  |
|  | 165 | 45 |  | 98.7 | 0.6 | 0.6 | 98.3 | 2.1 | 2.1 | -0.1 |  |
|  | 210 | 90 |  | 98.8 | 0.6 | 0.6 | 98.2 | 2.1 | 2.2 | -0.2 |  |
| Ximeixin 500 mg (batch number: 22303164) | 0 | N/A | Acid Stage ≤10% dissolved after 2 hours  Buffer Stage Q ≥75% within 45 min^a^ | 0.0 | 0 | N/A | 0.0 | 0 | N/A | N/A | N/A |
|  | 120 | 0 |  | 0.0 | 0.0 | 97.3 | 0.0 | 0.0 | 244.9 |  |  |
|  | 125 | 5 |  | 0.1 | 0.1 | 54.4 | 0.1 | 0.1 | 100.3 |  |  |
|  | 130 | 10 |  | 0.1 | 0.1 | 135.1 | 0.3 | 0.3 | 110.0 |  |  |
|  | 135 | 15 |  | 5.6 | 3.3 | 59.7 | 9.1 | 7.0 | 77.0 |  |  |
|  | 140 | 20 |  | 22.3 | 7.9 | 35.5 | 35.8 | 10.7 | 30.0 |  |  |
|  | 150 | 30 |  | 80.4 | 7.7 | 9.6 | 91.7 | 4.5 | 4.9 |  |  |
|  | 165 | 45 |  | 96.4 | 3.1 | 3.2 | 97.9 | 1.8 | 1.8 | 0.5 | -3.7 |
|  | 210 | 90 |  | 96.5 | 3.1 | 3.2 | 97.7 | 1.9 | 2.0 | 0.4 | -2.0 |
| Samelix 500 mg (batch number: 10323) | 0 | N/A | Acid Stage ≤10% dissolved after 2 hours  Buffer Stage Q ≥75% within 90 min^a^ | 0.0 | 0 | N/A | 0.0 | 0 | N/A | N/A | N/A |
|  | 120 | 0 |  | 0.0 | 0.0 | 60.0 | 0.0 | 0.1 | 157.5 |  |  |
|  | 125 | 5 |  | 0.1 | 0.1 | 65.5 | 0.2 | 0.1 | 78.5 |  |  |
|  | 130 | 10 |  | 0.2 | 0.1 | 24.4 | 0.2 | 0.1 | 58.5 |  |  |
|  | 135 | 15 |  | 0.9 | 0.5 | 55.8 | 2.2 | 1.2 | 52.4 |  |  |
|  | 140 | 20 |  | 5.9 | 4.7 | 80.7 | 30.1 | 12.8 | 42.6 |  |  |
|  | 150 | 30 |  | 73.2 | 13.5 | 18.4 | 94.5 | 4.7 | 5.0 |  |  |
|  | 165 | 45 |  | 96.9 | 1.7 | 1.8 | 96.5 | 1.2 | 1.2 | -0.1 | 1.0 |
|  | 210 | 90 |  | 96.9 | 1.7 | 1.8 | 96.7 | 1.2 | 1.2 | -0.1 | -0.2 |
| Hepamethion 500 mg (batch number: 75626) | 0 | N/A | Acid Stage ≤10% dissolved after 2 hours  Buffer Stage Q ≥75% within 90 min^a^ | 0.0 | 0 | N/A | 0.0 | 0 | N/A | N/A | N/A |
|  | 120 | 0 |  | 3.3 | 7.7 | 230.7 | 0.1 | 0.1 | 155.7 |  |  |
|  | 125 | 5 |  | 3.6 | 8.1 | 226.5 | 0.1 | 0.2 | 137.7 |  |  |
|  | 130 | 10 |  | 3.7 | 8.4 | 224.2 | 1.0 | 0.8 | 85.2 |  |  |
|  | 135 | 15 |  | 5.8 | 8.7 | 149.8 | 31.1 | 16.0 | 51.6 |  |  |
|  | 140 | 20 |  | 24.1 | 23.5 | 97.2 | 72.4 | 25.1 | 34.7 |  |  |
|  | 150 | 30 |  | 80.7 | 16.9 | 20.9 | 98.0 | 1.7 | 1.7 |  |  |
|  | 165 | 45 |  | 97.4 | 2.2 | 2.3 | 98.3 | 1.8 | 1.9 | 0.3 | -2.2 |
|  | 210 | 90 |  | 97.7 | 2.2 | 2.2 | 98.2 | 1.8 | 1.8 | 0.2 | -2.5 |
| Hepamethion 500 mg (batch number: 63551) | 0 | N/A | Acid Stage ≤10% dissolved after 2 hours  Buffer Stage Q ≥75% within 90 min^a^ | 0.0 | 0.0 | N/A | 0.0 | 0.0 | N/A | N/A | N/A |
|  | 120 | 0 |  | 0.1 | 0.2 | 244.9 | 0.1 | 0.1 | 165.7 |  |  |
|  | 125 | 5 |  | 0.1 | 0.3 | 244.9 | 0.1 | 0.2 | 231.3 |  |  |
|  | 130 | 10 |  | 0.5 | 1.0 | 211.8 | 0.8 | 1.6 | 213.4 |  |  |
|  | 135 | 15 |  | 6.7 | 8.0 | 119.2 | 13.3 | 13.0 | 98.2 |  |  |
|  | 140 | 20 |  | 27.6 | 20.4 | 73.9 | 42.1 | 14.6 | 34.8 |  |  |
|  | 150 | 30 |  | 74.1 | 35.9 | 48.4 | 93.9 | 4.0 | 4.2 |  |  |
|  | 165 | 45 |  | 95.6 | 5.7 | 6.0 | 98.5 | 1.9 | 1.9 | 1.0 | -7.2 |
|  | 210 | 90 |  | 98.4 | 3.1 | 3.2 | 98.5 | 1.8 | 1.8 | 0.0 | 0.2 |
| Hepamethion 500 mg (batch number: 69005) | 0 | N/A | Acid Stage ≤10% dissolved after 2 hours  Buffer Stage Q ≥75% within 90 min^a^ | 0.0 | 0.0 | N/A | No data^c^ | | | N/A | N/A |
|  | 120 | 0 |  | 14.6 | 35.3 | 241.7 |  |  |  |  |  |
|  | 125 | 5 |  | 14.5 | 34.9 | 240.8 |  |  |  |  |  |
|  | 130 | 10 |  | 14.6 | 35.0 | 239.0 |  |  |  |  |  |
|  | 135 | 15 |  | 18.3 | 36.5 | 199.2 |  |  |  |  |  |
|  | 140 | 20 |  | 41.5 | 34.2 | 82.4 |  |  |  |  |  |
|  | 150 | 30 |  | 92.2 | 12.8 | 13.9 |  |  |  |  |  |
|  | 165 | 45 |  | 99.3 | 1.5 | 1.5 |  |  |  |  |  |
|  | 210 | 90 |  | 99.3 | 1.5 | 1.5 |  |  |  |  |  |
| Adenomak 500 mg (batch number: KAY2301A) | 0 | N/A | Acid Stage ≤10% dissolved after 2 hours  Buffer Stage Q ≥75% within 90 min^a^ | 0.0 | 0.0 | N/A | No data^d^ | | | N/A | N/A |
|  | 120 | 0 |  | 0.0 | 0.0 | 152.3 |  |  |  |  |  |
|  | 125 | 5 |  | 0.0 | 0.0 | 244.9 |  |  |  |  |  |
|  | 130 | 10 |  | 0.0 | 0.0 | 244.9 |  |  |  |  |  |
|  | 135 | 15 |  | 0.3 | 0.1 | 39.0 |  |  |  |  |  |
|  | 140 | 20 |  | 1.6 | 1.0 | 65.0 |  |  |  |  |  |
|  | 150 | 30 |  | 9.0 | 3.1 | 34.4 |  |  |  |  |  |
|  | 165 | 45 |  | 19.7 | 6.9 | 35.3 |  |  |  |  |  |
|  | 210 | 90 |  | 57.5 | 28.7 | 49.8 |  |  |  |  |  |
| Adenomak 500 mg (batch number: KAY2219A) | 0 | N/A | Acid Stage ≤10% dissolved after 2 hours  Buffer Stage Q ≥75% within 90 min^a^ | 0.0 | 0.0 | N/A | No data^d^ | | | N/A | N/A |
|  | 120 | 0 |  | 0.1 | 0.2 | 119.7 |  |  |  |  |  |
|  | 125 | 5 |  | 0.3 | 0.2 | 74.7 |  |  |  |  |  |
|  | 130 | 10 |  | 0.3 | 0.2 | 63.7 |  |  |  |  |  |
|  | 135 | 15 |  | 0.6 | 0.3 | 53.1 |  |  |  |  |  |
|  | 140 | 20 |  | 1.9 | 1.1 | 56.9 |  |  |  |  |  |
|  | 150 | 30 |  | 9.8 | 1.2 | 11.8 |  |  |  |  |  |
|  | 165 | 45 |  | 24.4 | 6.0 | 24.5 |  |  |  |  |  |
|  | 210 | 90 |  | 75.0 | 21.9 | 29.2 |  |  |  |  |  |
| Adenomak 500 mg (batch number: KAY2225A) | 0 | N/A | Acid Stage ≤10% dissolved after 2 hours  Buffer Stage Q ≥75% within 90 min^a^ | 0.0 | 0.0 | N/A | No data^d^ | | | N/A | N/A |
|  | 120 | 0 |  | 0.0 | 0.0 | N/A |  |  |  |  |  |
|  | 125 | 5 |  | 0.0 | 0.0 | 244.9 |  |  |  |  |  |
|  | 130 | 10 |  | 0.1 | 0.0 | 84.1 |  |  |  |  |  |
|  | 135 | 15 |  | 0.2 | 0.2 | 77.2 |  |  |  |  |  |
|  | 140 | 20 |  | 0.6 | 0.4 | 73.7 |  |  |  |  |  |
|  | 150 | 30 |  | 2.5 | 1.8 | 73.1 |  |  |  |  |  |
|  | 165 | 45 |  | 12.4 | 5.2 | 42.2 |  |  |  |  |  |
|  | 210 | 90 |  | 39.6 | 17.5 | 44.1 |  |  |  |  |  |

^a^Dissolution test for solid dosage forms, delayed release dosage forms level B1. No value is less than Q=5%; ^b^Ratio = stability slope competitor / stability slope Heptral. Similar stability slope results in a ratio equal to 1; ^c^Dissolution run interrupted due to instrument failure. Due to limited availability, no additional tablets available to repeat the analysis; ^d^After being exposed to the dissolution media, the tablets became adhesive and stuck to the dissolution vessel. No disintegration visible after 210 min / 90 min in buffer stage.

CV, coefficient of variation; N/A, not applicable; RH, relative humidity; SD, standard deviation; T=0, Month 0, initial analysis; T=3M, Month.
